# Cortical dynamics following real-time MEG neurofeedback training of the speed of shifting spatial attention: a pilot study

**DOI:** 10.1101/2022.06.19.496738

**Authors:** Kunjan D. Rana, Sheraz Khan, Matti S. Hämäläinen, Lucia M. Vaina

## Abstract

Neurofeedback is a technique that directs self-regulated modulation of neural activity. This is achieved by delivering real-time feedback derived from brain signals. In our previous work, we introduced a novel method, state-based neurofeedback (sb-NFB) that targets patterns of MEG signals related to shifts in spatial attention. In this pilot study, we used sb-NFB to train participants to decrease the time required to covertly shift spatial attention from one visual hemifield to the other. We characterized the changes to cortical connectivity during each training session. In addition, we run a separate, Posner-like validation task before the training sessions and after the training was complete. We found a significant main effect of training on the reaction time linked to switching spatial attention in both the training task and the validation task. This indicates the achieved improvement in shifting spatial attention generalized to another situation requiring this capability.

## 1. INTRODUCTION

Recent studies demonstrate that MEG can be successfully used as a real-time neurofeedback device, that allows subjects to modulate and enhance task-related cortical rhythms associated with sensory, motor, or cognitive performance in both healthy subjects and clinical populations (Bagherzadeh et al., 2020; Foldes et al., 2011; Foldes et al., 2015; Okazaki et al., 2015; Parkkonen, 2015). Subjects undergoing neurofeedback training acquire a strategy to self-regulate brain activity either through conditioning or through their own volition. By receiving sensory feedback related to brain activity, subjects learn how to control their brain activity through reinforcement learning. The purpose of the training is to control brain functions related to the measured cortical activity.

In our previous work, we have introduced and described the method of state-based neurofeedback (sb-NFB), which trains the time involved in changes between brain states, rather than the activation patterns themselves (Rana et al., 2020). We demonstrated via an online analysis that the sb-NFB method captures information about brain states by measuring oscillatory activity across all sensors in MEG. We used dimensionality reduction techniques to weight sensors and oscillatory frequency bands that optimally separate the targeted cognitive states. Linear support vector machines (SVMs) were applied to decode a cognitive state in real-time from the dimensionally reduced dataset.

In this study, we applied the sb-NFB method to train subjects to shorten the time necessary for shifting covert spatial attention from one visual hemifield to the other through an offline analysis. We refer to this time as the “switch time.” Here we uncover how neural dynamics evolve through changes in the cortical functional connectivity during the course of sb-NFB training. Furthermore, we show that sb-NFB training can generalize to other situations that require fast switching of spatial attention by tracking changes in the behavioral performance in a separate, unrelated spatial attention validation task (a Posner task).

## 2. MATERIALS AND METHODS

### 2.1 Participants

Seven subjects (mean age = 24.3, SD = 5.1, 4 females; all right-handed) participated in this study. They were all naïve to the purpose of the experiment. All participants had a normal or corrected-to-normal vision and had no known neurological or psychiatric conditions. Each participant signed a consent form according to a protocol approved by the Institutional Review Board of Boston University and Massachusetts Institute of Technology (MIT).

### 2.2 Overview of Training

Figure 1 shows the overview of the training sessions. As shown, on each of the six training days we employ the *NFB task* protocol whose purpose is to shorten the time required to switch spatial attention from one visual field to another. The NFB task is described in detail in Section 2.3 below. In addition, on the day before the first training session and the day after the last one, we administered an additional *validation task*, whose purpose is to find whether the achieved improvement in shifting spatial attention generalizes to another situation requiring the same capability. The validation task is described in Section 2.4. As indicated in Figure 1 there is one day of rest after three training days. The total length of the study for each participant is thus nine days.

**Figure 1.**
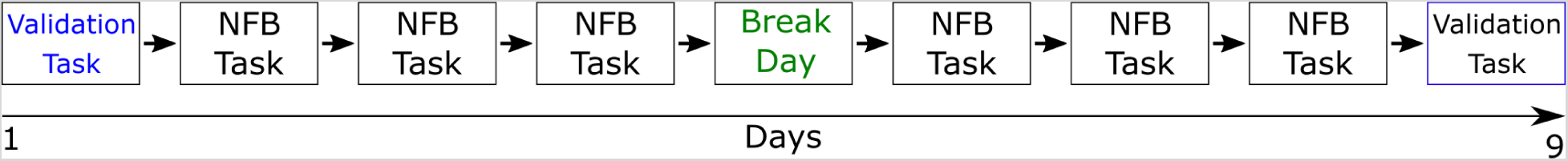
Overview of training. The diagram shows the sequence of training and validation sessions.

### 2.3 The NFB Task

The timing diagram of each of the NFB Task trials is shown in Figure 2. Each training session consisted of five blocks of 80 trials each. Figure 2 shows schematically the sequence of screens presented in chronological order. At the center of the screen, we show the feedback cue as a “thermometer”. The exact details of the stimulus parameters are described in (Rana et al., 2020).

**Figure 2.**
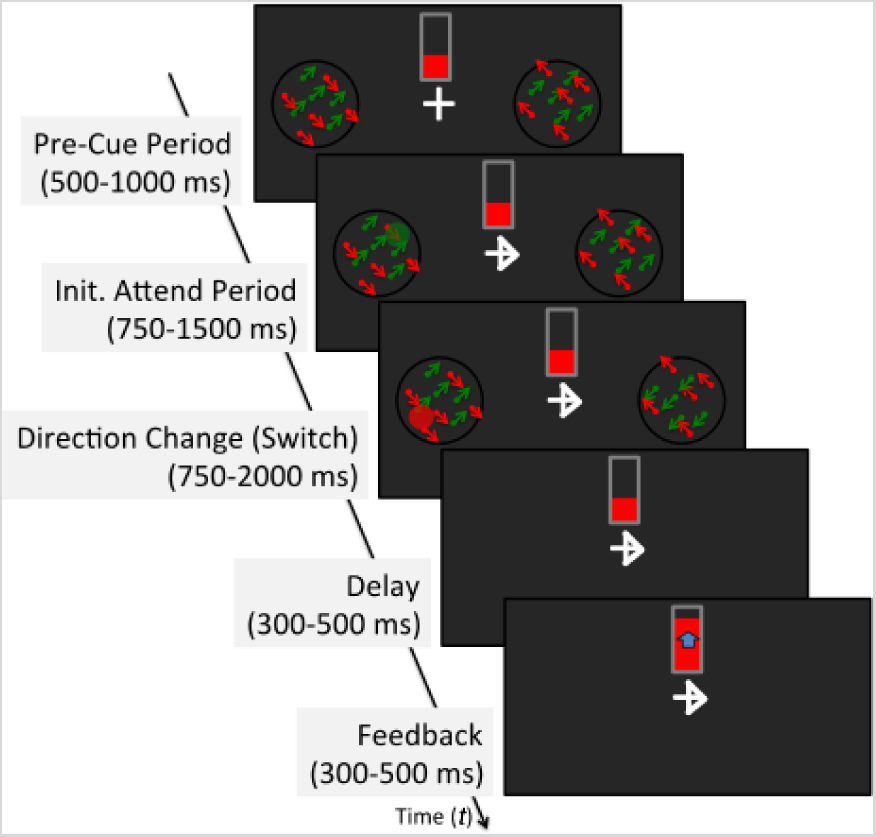
NFB task. At the beginning of each trial, a fixation mark is shown at the center of the screen. Participants are asked to maintain fixation on the fixation mark for the full duration of each trial. Above the fixation mark, there is a feedback thermometer, with height (indicated as the red bar) representing the speed of attention switch in the previous trial. On the left and right sides of the screen are two apertures equidistant from the fixation mark. Each aperture contains a random-dot kinematogram (RDK) consisting of an equal percentage of red and green dots. The different color dots in each aperture move with consistent diagonal planar motion, but each color moves orthogonally to the other. During the “Init. Attend Period”, the fixation cross is replaced with an arrow, which indicates the hemifield that the participant is required to attend. At this time, a disc is displayed in the opposite aperture and is set to change location and color (red or green) at random intervals. After a variable period of time, the dot set (green or red) in the attended may changes direction of motion; if the attended dots change direction, the participant switches attention to the aperture in the opposite hemifield. The two apertures are removed from the screen concurrently with the participant responding to the color of the disc in the unattended aperture. The feedback thermometer is then updated to reflect the switch time of the previously completed trial. Modified from (Rana et al., 2020).

The height of the thermometer was based on *z_t_* = (*ρ* − *µ_ρ_*)/*σ_ρ_*, the *z*-scored switch time, where *µ_ρ_* is the mean switch time and *σ_ρ_* is the standard deviation of switch times from the previous block of trials. The switch time *ρ* is derived from the brain state as explained in detail in (Rana et al., 2020).

### 2.4 The Validation Task

The timing diagram of each of the trials in the behavioral validation task is shown in Figure 3. The task consisted of three blocks of 80 trials. Each trial consisted of fixation, cue, and target presentations. The target, a rotating random-dot kinematogram (RDK), appears in the visual hemifield indicated by the arrow in 80% of the trials (valid cue), and on the opposite side in 20% of the trials (invalid cue). Regardless of the visual field location of the RDK, the participant is instructed to indicate the direction of dot motion (clockwise or counterclockwise) via a button box.

**Figure 3.**
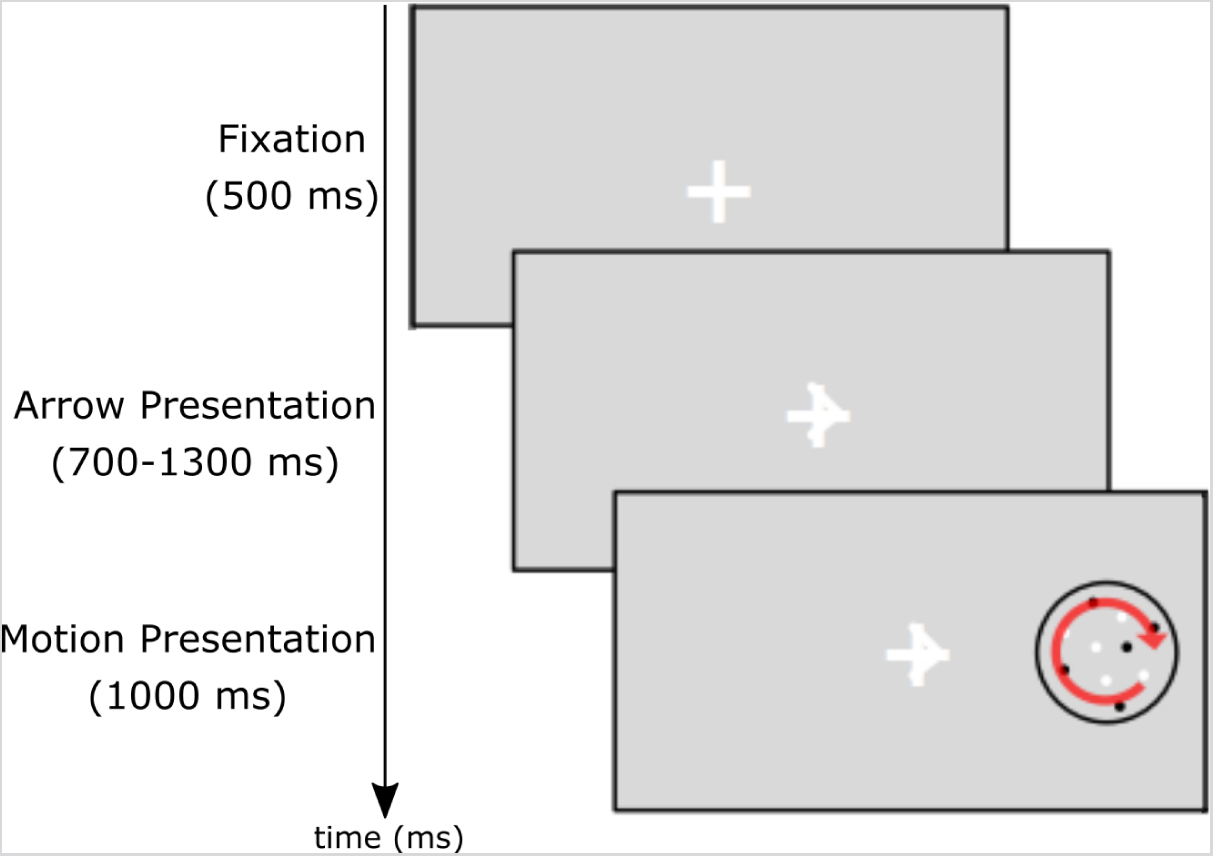
Spatial attention validation task. Each trial begins with a white fixation cross displayed at the center of an otherwise gray screen. Participants are instructed to fixate their gaze on the center of the screen for the full duration of each trial. After 500 ms the fixation cross changes into an arrow pointing either to the left or to the right. The participant is instructed to attend to the visual hemifield pointed to by the arrow. After a variable period of 700 – 1300 ms, a gray RDK pattern appears, with dots moving clockwise or counterclockwise. The RDK pattern consisted of a circular aperture subtending 4° in diameter (gray dots, density 4 dots/deg^2^; diameter 0.1°; luminance 40 cd/m^2^; background luminance 25 cd/m^2^, motion velocity 3 rad/sec). In the motion aperture, 90% of the dots provided a motion signal and moved either clockwise or counterclockwise. The rest of 10% of the dots provided motion noise. The center of the aperture was positioned horizontally at 8° to the left or right from the central fixation cross. The background luminance was 25 cd/m^2^. The gray fixation cross and the arrow were luminance 40 cd/m2 and were fit in a virtual square 0.5° in width and height.

### 2.5 MEG Data Acquisition

The magnetoencephalography (MEG) study was conducted at the Athinoula A. Martinos Imaging Center at MIT’s McGovern Institute for Brain Research. Participants were seated in a chair under the MEG sensor array in a three-layer magnetically shielded room (Ak3b, Vacuumschmelze GmbH, Hanau, Germany) and faced a projection screen placed at a distance of 138 cm. The stimuli were projected onto a 44” screen through an aperture in the MEG chamber using a Panasonic DLP projector (Model #PT-D7500U). During the experiment, the room lighting was dimmed. The MEG data were acquired with a 306-channel Neuromag Vectorview whole-head system (MEGIN OY, Helsinki, Finland), comprising of 204 planar gradiometers and 102 magnetometers.

To measure the participant’s head position during the MEG recordings, five head-position indicator (HPI) coils were fixed on the participant’s head. The positions of the HPI electrodes and at least 80 points on the scalp were digitized (Fastrak digitizer, Polhemus Inc., Colchester, VT) in a head coordinate frame defined by the nasion and the left and right auricular points for subsequent alignment with the anatomical MRI data. In the off-line analysis all data were aligned to a common head position through the Maxfilter software (Taulu et al., 2004; Taulu and Simola, 2006; Taulu et al., 2005).

Participants were instructed to avoid blinking during trials and blink only during the interstimulus intervals.

### 2.6 Anatomical MRI Acquisition

High-resolution T1 weighted structural Magnetic resonance images (MRI) were acquired on a separate day using an 8-channel phased-array head coil in a 3T scanner (Siemens-Trio, Erlangen, Germany). Parameters of the sequence were as follows: distance factor: 50%, slices per slab: 128, FOV: 256 mm, FOV phase: 100 degrees, slice thickness: 1.33 mm, TR: 2530 ms, TE: 3.39 ms.

### 2.7 MEG Data Preprocessing and Source Estimation

In the NFB task, MEG sensor data were segmented into separate trials from -1000 ms to 3000 ms relative to the onset of motion direction change in the attended aperture. Trials were rejected if the peak-to-peak amplitude exceeded 3000 fT and 3000 fT/cm in any of the magnetometer and gradiometer channels. To remove line noise artifacts, the MEG signals were notch filtered at 60, 120, and 180 Hz.

Freesurfer was used to construct the cortical 3D surface (Fischl et al., 2002a; Fischl et al., 2002b; Fischl et al., 1999b; Fischl et al., 2004a; Fischl et al., 2004b) with approximately 300,000 vertices. Surfaces were decimated to 8000 vertices per hemisphere. Using the Brainstorm software (Tadel et al., 2011), the MEG sensor signals were mapped onto the reconstructed 3D cortical surface. Brainstorm was also used to compute the forward gain matrix overlapping spheres model which maps current strengths on the cortex to MEG signal amplitudes recorded by the sensor array (Huang et al., 1999). We constrained the current orientation to be normal to the cortical manifold. We used the minimum norm estimate (MNE) method (Hämäläinen and Ilmoniemi, 1994) to compute the source estimate, which maps the sensor measurements back onto the cortical surface. For this computation, the noise-covariance matrix was estimated from a 500 ms window immediately before the pre-cue period. We visualized the result as a dynamic statistical parametric map (Dale et al., 2000), obtained by normalizing the activity by the noise estimate mapped to the cortical surface.

### 2.8 Cortical MEG measures

Based on source estimates, we defined clusters of activation on the cortex and used them as a basis for defining the regions of interest (ROI) shown in Figure 4. To obtain a common set of regions for all participants, we used Freesurfer to morph the dSPM from the individual participants’ cortical surface onto the FsAverage (Fischl et al., 1999a; Fischl et al., 1999b). The ROIs were defined by those clusters whose average activation during the *Attend window* (see Section 2.10) was significantly higher than the average activity over the interstimulus interval (z > 2) and were common (with overlap) to all the participants. MT+ (Calabro and Vaina, 2012; Martinez-Trujillo et al., 2007) and VIP (Calabro and Vaina, 2012; Field and Wann, 2005) labels were assigned based on literature-defined functional areas with a center of mass in the range of Talairach coordinates of the same areas defined in fMRI studies.

**Figure 4.**
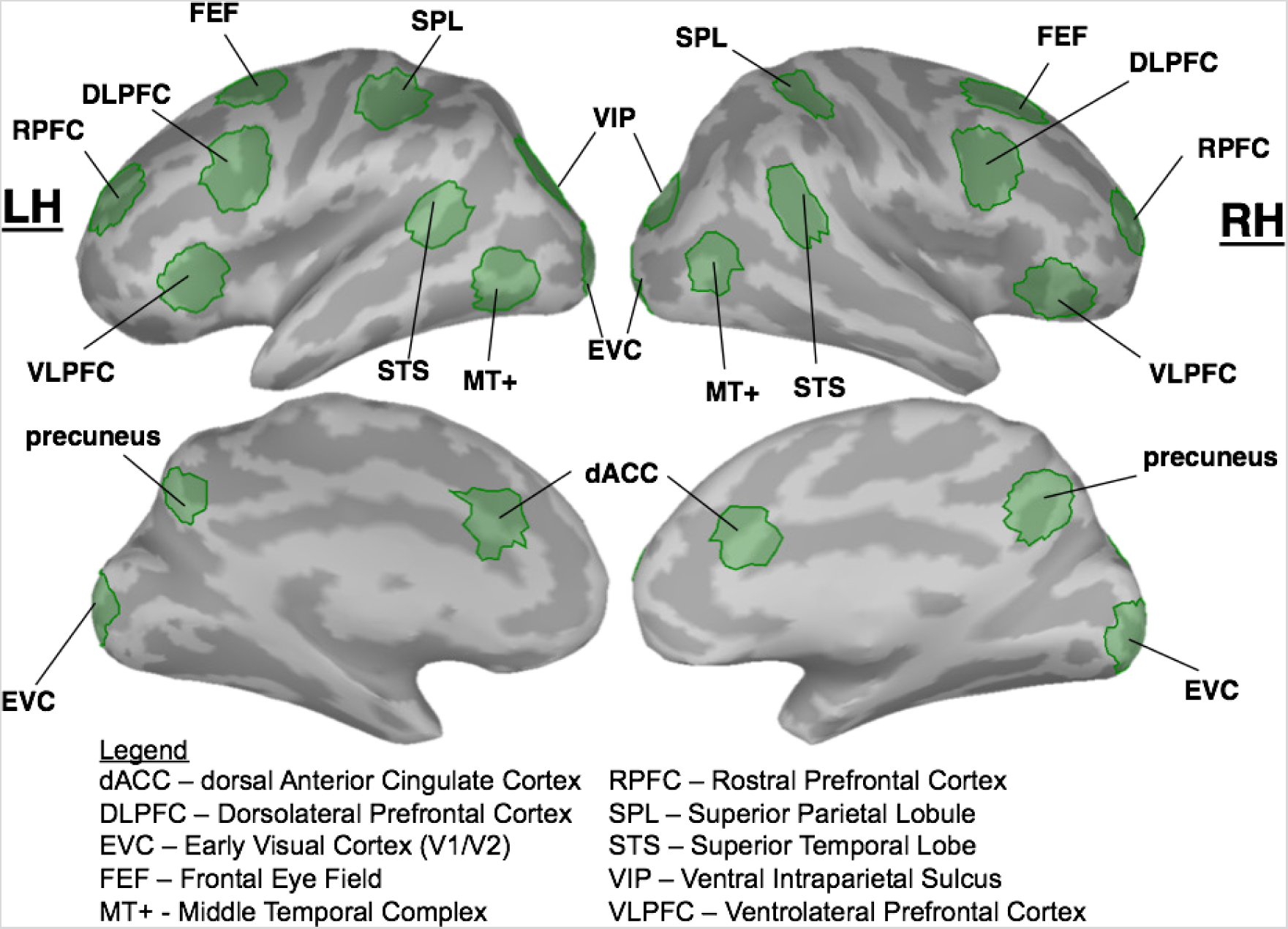
Functionally defined regions of interests (ROIs)

### 2.9 Cortical Functional Connectivity

The directional functional connectivity was quantified by Frequency-Domain Granger Causality (FDGC), computed using the Multivariate Granger Causality (MVGC) Toolbox (Barnett and Seth, 2014) and default parameters. FDGC is the spectral version of Granger Causality (Geweke, 1982). The frequency bands used for the FDGC analysis were alpha (8 – 13 Hz), beta (13 – 30 Hz), and gamma (30 – 60 Hz). To obtain a representative score for each frequency band, the Granger scores were average across each frequency band.

### 2.10 Analysis of the sb-Task

In the analysis of the sb-NFB task (Figure 2), we considered the following two windows as these were directly involved in the NFB training.

(i) The *Attend window*: from -500 to 0 ms with respect to the time of change in the motion direction in the RDK, *t_D_*.
(ii) The *Switch window* from the motion direction change to the average time required to switch attention, *t*_switch_.

To find connections that changed in Granger score consistently across participants over the 6 training days, we first computed FDGC in each of the two time windows. In the “attend” window, participants prepared to switch attention when the motion direction changed, and in the “switch” window, they switched spatial attention from the attended aperture to the aperture in the opposite visual hemifield.

We performed mixed ANOVA analysis for each connection (Matlab function ANOVAN) within the time windows in each frequency band with the training session as a within-subject factor and the participant ID as a between-subjects factor. If a connection was deemed to be significant for the within-subject (p < 0.05) factor, we computed posthoc Pearson correlation on the averaged GC score (across subjects) to determine whether the connection was increasing or decreasing.

## 3. RESULTS

We organize our results to describe: (i) Changes in cortical functional connectivity during sb-NFB training, and (ii) Changes in reaction times in the validation task.

### 3.1 Changes in Functional Connectivity during sb-NFB Training

The sustained changes in functional connectivity across all training sessions in the attend and switch time windows are summarized in Figure 5. The connections between the ROIs that increased or decreased significantly in Granger score during the sb-NFB training are shown in Figure 5A. We report the connections that had a significant effect on the training day factor. An increase in Granger score *A* → *B* suggests that the causal influence of *A* to *B* increases and vice versa. This kind of directional influence is often called directional functional connectivity. The number of increasing and decreasing connections is summarized in Figure 5B. During the attend window in the alpha band, a larger number of connections decreased (n = 10) in comparison to the number of connections increased (n = 6). In the beta and gamma bands, all connections increased in strength during training. In the switch window, a larger number of connections in the gamma band increased (n = 32) than in the attend window (n = 16). Overall, connectivity increased over the course of training, with the gamma band having the most pronounced increase.

**Figure 5.**
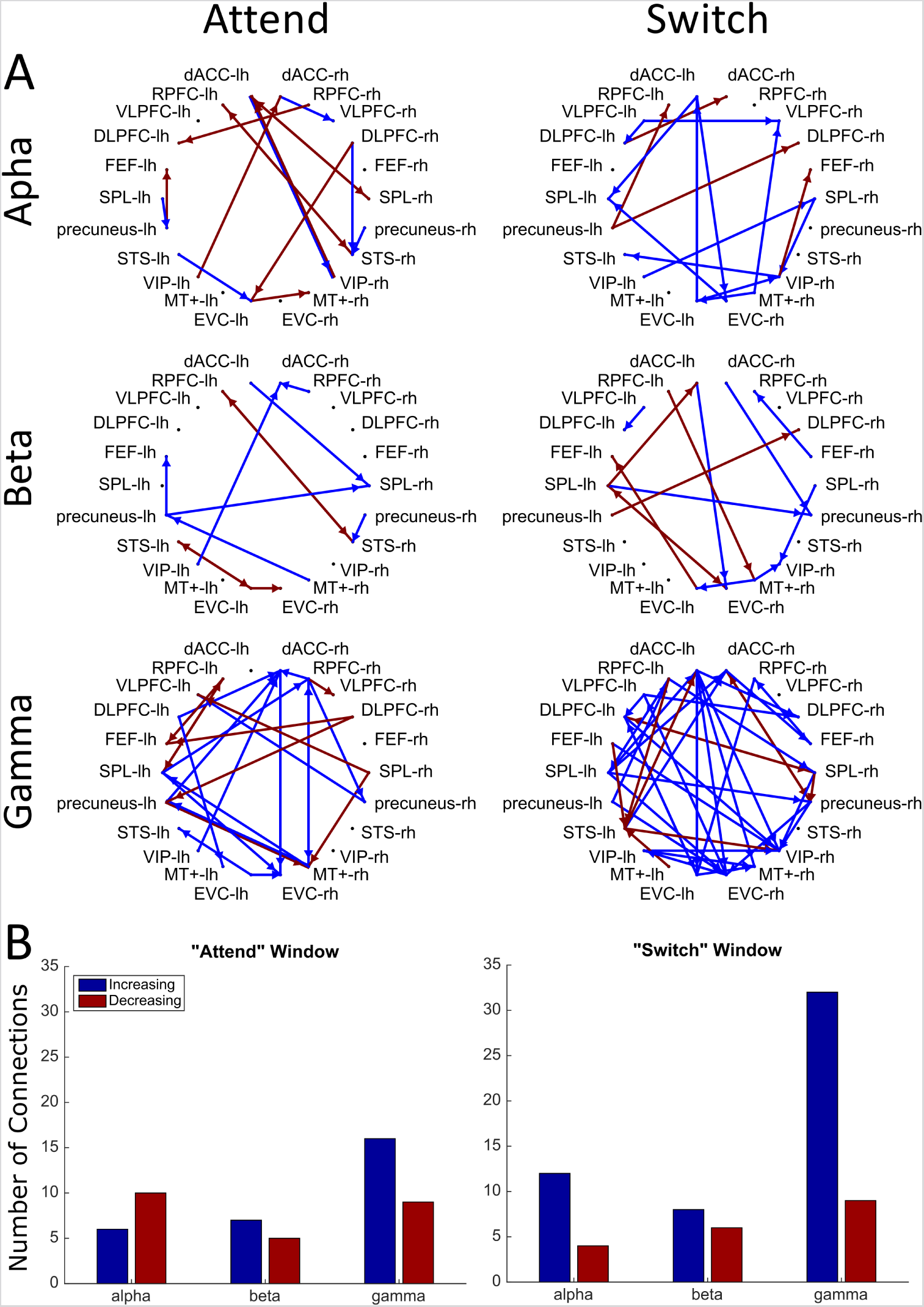
Changes in directed functional connectivity in the attend and switch time windows during sb-NFB training. **(A)** Each circle plot shows connections showing systematic evolution during the training days (ANOVA, p < 0.05) with blue lines representing increases and red lines representing decreases over the course of NFB training, respectively. The top row represents alpha-band, the middle row beta-band, and the bottom row gamma-band connectivity in the attend (left) and switch (right) windows. The ROIs are labeled as dACC – dorsal Anterior Cingulate Cortex, DLPFC – Dorsolateral Prefrontal Cortex, EVC – Early Visual Cortex, FEF – Frontal Eye Field, MT+ – Middle Temporal Cortex, RPFC – Rostral Prefrontal Cortex, SPL – Superior Parietal Lobe, STS – Superior Temporal Lobe, VIP – Ventral Intraparietal Sulcus, VLPFC – Ventrolateral Prefrontal Cortex. **(B)** The number of connections that increased (blue) or decreased (red) during NFB training in the attend (left) and switch windows (right) in the alpha, beta, and gamma bands. The height of each bar is the count of the connections that increased or decreased in strength over the training period.

### 3.2. Changes in Reaction Times in the Validation Task (Transfer Learning)

To ensure the generalizability of the neurofeedback training, we measured behavioral performance in the spatial attention validation task before and after the sb-NFB training for each subject. Even before NFB training, the percent correct in the validation task across all participants was at the ceiling (98.09% mean), while reaction time was more variable (691.7 +/- 233.4 ms). Therefore, we focused the behavioral analysis on the reaction time.

Figure 6 illustrates the mean and standard errors of the day and cue validity factors with respect to the reaction time. There was a significant interaction with the day and the cue validity (F (1,3325) = 4.325, *p* = 0.038), as well as significant main effects of the day (F(1,3325) = 180.194, *p* < 0.001) and the cue validity (F(1,3325) = 180.194, *p* < 0.001). There was a significant interaction between cue validity and attended location (F (1,3325) = 19.669, *p* < 0.001), but no interaction with the day (before or after NFB training) suggesting that participants were biased in responding to motion in one hemifield (either left or right).

**Figure 6.**
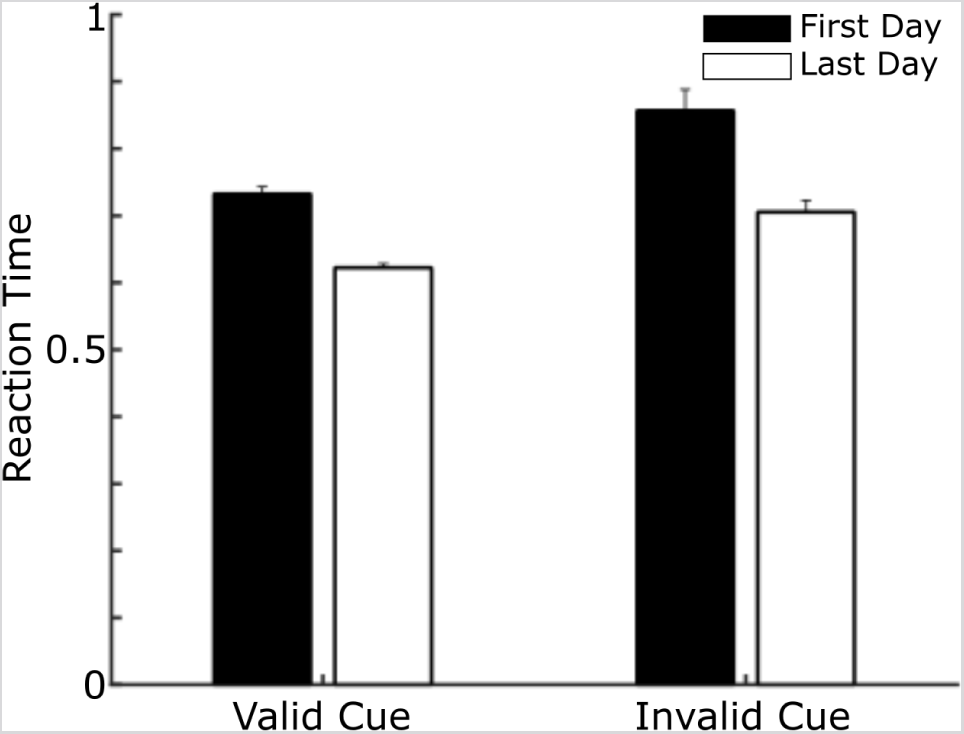
Reaction time in spatial attention validation task by cue validity (i.e., valid or invalid) and day (i.e., first or last day). Error bar represents one standard error above the mean reaction time. The y-axis gives the reaction time in seconds.

## 4. DISCUSSION

This paper describes the functional changes in brain connectivity using the sb-NFB method (Rana et al., 2020). Our results suggest that the sb-NFB method alters neural connectivity across frequency bands during the attend as well as the switch window.

In the attend window, subjects attended to the visual hemifield where they were cued to expect the motion direction switch. We suggest that the strengthened beta-band connections over sb-NFB training may be due to increased attentiveness to the motion direction change. Since the beta-band is implicated in top-down attention (Bastos et al., 2015; Khan et al., 2018) connectivity to MT+ suggests that sustained attention is directed towards the motion of the cued stimulus to detect the change in direction of motion. The subjects that had greater improvement on the speed of changing attention from one visual field to the other had higher sustained attention to the cue.

### 4.1 sb-NFB Training Results in Weaker Signals during Targeted Behavior

There was a significant increase in the strength of gamma-band connectivity amongst RPFC, VLPFC, FEF, and SPL ROIs during the “switch”. However, in other cortical regions, the strength of gamma connectivity was lower. We suspect that more efficient engagement is the reason for the lower gamma connectivity strength. The gamma-band is shown to be involved in bottom-up sensory processing (Bastos et al., 2015; Khan et al., 2018; Kinreich et al., 2017) has been suggested to be a physiological fingerprint of attention (Jensen et al., 2007). Since the “switch” time decreases with training days, the network implementing the “switch” must also last for a shorter duration.

### 4.2 Transfer Learning to a Spatial Attention Validation Task

To assess whether there was a transfer of learning a faster switch speed we administered a spatial attention validation task before and after subjects were administered the training protocol. The spatial attention control task is a Posner-like cueing task, in which subjects may be required to change their spatial attention. Throughout this task, the subjects fixated on a centrally presented cross. At the beginning of each trial, subjects were cued to a visual field (left or right) and then asked to respond to the direction of motion of an RDK pattern that appeared after a random interval of time (validly-cued trials). In 20% of trials, the RDK pattern appeared on the opposite side (invalidly-cued trials). In these trials, participants were required to switch attention to the opposite visual field and respond to the direction of motion in the RDK pattern on the opposite side.

Our results show that after the completion of the NFB training, there was a significant decrease in the time required to shift spatial attention for both validly- and invalidly-cued trials. This suggests that sb-NFB training helped participants to successfully increase their speed of switching spatial attention, resulting in a reduced response time when attention switching was necessary.

## 5. CONCLUSION AND LIMITATIONS

In this paper, we showed that the sb-NFB training paradigm leads to changes in functional connectivity and these changes generalize to similar situations. Previous MEG and EEG neurofeedback studies monitored changes to activation within a particular trained sensor or ROI (Bagherzadeh et al., 2020; Hammond, 2003; Lansbergen et al., 2011; Okazaki et al., 2015; Sudre et al., 2011; Wang et al., 2016). However, since we trained on the dynamics of “brain state” using all sensor measures, we expected large-scale changes to cortical connectivity.

We suggest that the application of the sb-NFB method illustrated here and our findings may provide a framework for further studies in both healthy and neurological subjects, such as patients with hemispatial neglect and other spatial attention deficits. To measure whether sb-NFB training is retained, we would need to re-test healthy and clinical participants at some time interval after the training protocol has ended. Furthermore, we will have to demonstrate that improvement is due to NFB training. For instance, a sham training indicator could be used to compare against sb-NFB training. In addition, our task in which we measured transfer of learning involved a cue that was a motion stimulus. Although this study reveals that there is a transfer of learning onto this task, we cannot delineate the changes between detection of the motion stimulus and the switching ability, which both would result in faster switching time. Changing the cue type to a non-motion cue (e.g. color, shape) would help identify the transferred learning effect. Our results suggest several changes to connectivity patterns after training, demonstrating that sb-NFB does indeed lead to changes in functional connectivity. With more data, we expect to be able to finely delineate the connectivity changes.

